# Acoustic tumor paint for real-time imaging, surgical guidance and recurrence monitoring of brain tumors with ultrasound

**DOI:** 10.1101/2024.12.22.629782

**Authors:** Claire Rabut, George H. Daghlian, Pierina Barturen-Larrea, Hongyi Richard Li, Ruth Vorder Bruegge, Rebecca M. Jones, Dina Malounda, Gianmarco F. Pinton, Mikhail G. Shapiro

**Author notes:** These authors contributed equally to this work (Claire Rabut, George H. Daghlian). Corresponding author (MGS).

## Abstract

The rapid growth, invasiveness, and resistance to treatment of glioblastoma multiforme (GBM) underscore the urgent need for improved diagnostics and therapies. Current surgical practice is limited by challenges with intraoperative imaging, while recurrence monitoring requires expensive magnetic resonance or nuclear imaging scans. Here we introduce “acoustic tumor paint”, an approach to labeling brain tumors for ultrasound imaging – a widely accessible imaging modality. We show that gas vesicles (GVs), natural air-filled protein nanostructures, preferentially accumulate in brain tumors following systemic administration in syngeneic and xenograft mouse models of GBM. This enables real-time tumor visualization during surgery and postoperative monitoring of recurrence. We characterize GV uptake and breakdown by tumors and their resident cells and support clinical translatability by documenting non-toxic repeated administration. We also demonstrate the potential for post-operative monitoring in humans by imaging GVs through a human skull and an FDA-approved skull prosthesis. Acoustic tumor paint has the potential to enhance diagnostic accuracy, improve surgical outcomes, make monitoring more accessible, and extend survival in GBM patients.

## INTRODUCTION

Glioblastoma multiforme (GBM) poses a substantial challenge within the field of neuro-oncology as the most common malignant primary brain tumor^1^. GBM is characterized by a high mortality rate and poor clinical outcomes, with a median survival time of 12 to 15 months after diagnosis^2,3^. This grim prognosis stems from the tumor’s rapid rate of cellular proliferation, extensive infiltration into neighboring healthy tissue, resistance to conventional therapeutic modalities, and a predisposition for rapid recurrence. GBM typically recurs in 6 to 9 months^4^.

Treatment for GBM consists of combined chemoradiotherapy and surgical resection^5,6^, aiming to balance maximal bulk tumor removal against potential neurological complications due to the removal of functional brain tissue^7^. Following detailed neurological evaluation, contrast-enhanced magnetic resonance imaging (MRI) is commonly used for initial tumor imaging and surgical planning, then repeated for postoperative monitoring of recurrence. Even with advances in neuronavigation relying upon the patient’s MRI scans and intraoperative optical labels such as sodium fluorescein and 5-aminolevulinic acid^8,9^, there are limitations to precisely resecting the tumor. These range from intraoperative brain shift (which introduces neuronavigation errors) to the inherent limitations of optical imaging methods at depths >1 mm due to light scattering and attenuation^9–15^. Furthermore, patients who live in resource-constrained areas or are too clinically unstable to undergo lengthy imaging sessions have limited access to MRI, making recurrence monitoring more difficult. In addition, there are unresolved concerns about the long-term safety of gadolinium-based MRI contrast agents^16,17^.

What if instead of MRI or optical imaging, brain tumors could be visualized with ultrasound, a versatile and inexpensive point-of-care imaging modality? Ultrasound offers excellent spatial and temporal resolution (order of 100 µm and 1 ms), provides centimeter scale tissue penetration, and has extensively documented clinical safety^18,19^. Although ultrasound is already utilized in neurosurgical procedures, no contrast agent currently exists to label brain tumors specifically for this modality^11,12,20^. We hypothesized that if we could equip ultrasound with a suitable tumor-specific contrast agent, this would provide transformative capabilities for real-time surgical guidance and timely detection of tumor recurrence.

To develop a suitable brain-tumor label, we reasoned that a sufficiently small acoustic contrast agent could enter tumors through the disrupted blood-brain barrier (BBB) distinguishing them from healthy brain tissue^21^. This compromised and leaky BBB is already leveraged in the clinic for gadolinium-enhanced MRI and in emerging nanoparticle delivery approaches^22,23^. We postulated that an ideal “acoustic tumor paint” would be nanoscale (smallest dimension < 100 nm), selectively detectable with nonlinear ultrasound modes over background tissues, capable of sustained real-time imaging during surgery, and biodegradable to allow safe repeated administration.

These criteria led us to focus on gas vesicles (GVs), a recently introduced class of contrast agents for ultrasound imaging and therapy. These air-filled protein nanostructures are derived from certain aquatic microorganisms, which use them for buoyancy regulation^24,25^. The GVs used in this study are biconical tube-shaped structures measuring approximately 85 nm by 500 nm. Their 2 nm-thick protein shell prevents liquid water from entering the GV while allowing the dynamic exchange of gas with surrounding media, thus forming a stable nanoscale air pocket^26^. Acoustic waves are strongly scattered at this air–water interface, enabling GVs to produce robust ultrasound contrast when injected into the body or expressed in engineered cells^27–32^. Critically for in vivo imaging, GVs exhibit strongly nonlinear responses to acoustic pressure, which results in nonlinear scattering of ultrasound and enables effective differentiation from surrounding tissue^33–37^. Additionally, GVs are resilient to repeated insonification, amenable to molecular engineering to bind to specific markers or respond to biological functions, and have growing applications in therapeutic ultrasound, optical imaging, and MRI^28,38–42^. Prior work has suggested that GVs could accumulate in subcutaneous tumors in mice^29,43–45^, but they have not been tested in brain tumors or applied to surgical guidance or recurrence monitoring.

In this study, we show that, upon intravenous (IV) injection, GVs naturally and preferentially accumulate in brain tumor parenchyma, where they can be visualized with nonlinear ultrasound, effectively ‘painting’ the tumors acoustically before being degraded after approximately 1 hour (**Fig. 1a-b**). We confirm the generalizability of this labeling across three syngeneic and xenograft mouse models of GBM and characterize the mechanisms of GV accumulation and degradation using tissue histology. We demonstrate the use of acoustic tumor paint in ultrasound-guided surgery (**Fig. 1c**) and its ability to reveal recurrent tumors following resection. Finally, we establish that GVs are non-toxic with repeated administration and detectable through both the human skull and FDA-approved human skull implants, laying the groundwork for clinical translation.

**Figure 1:**
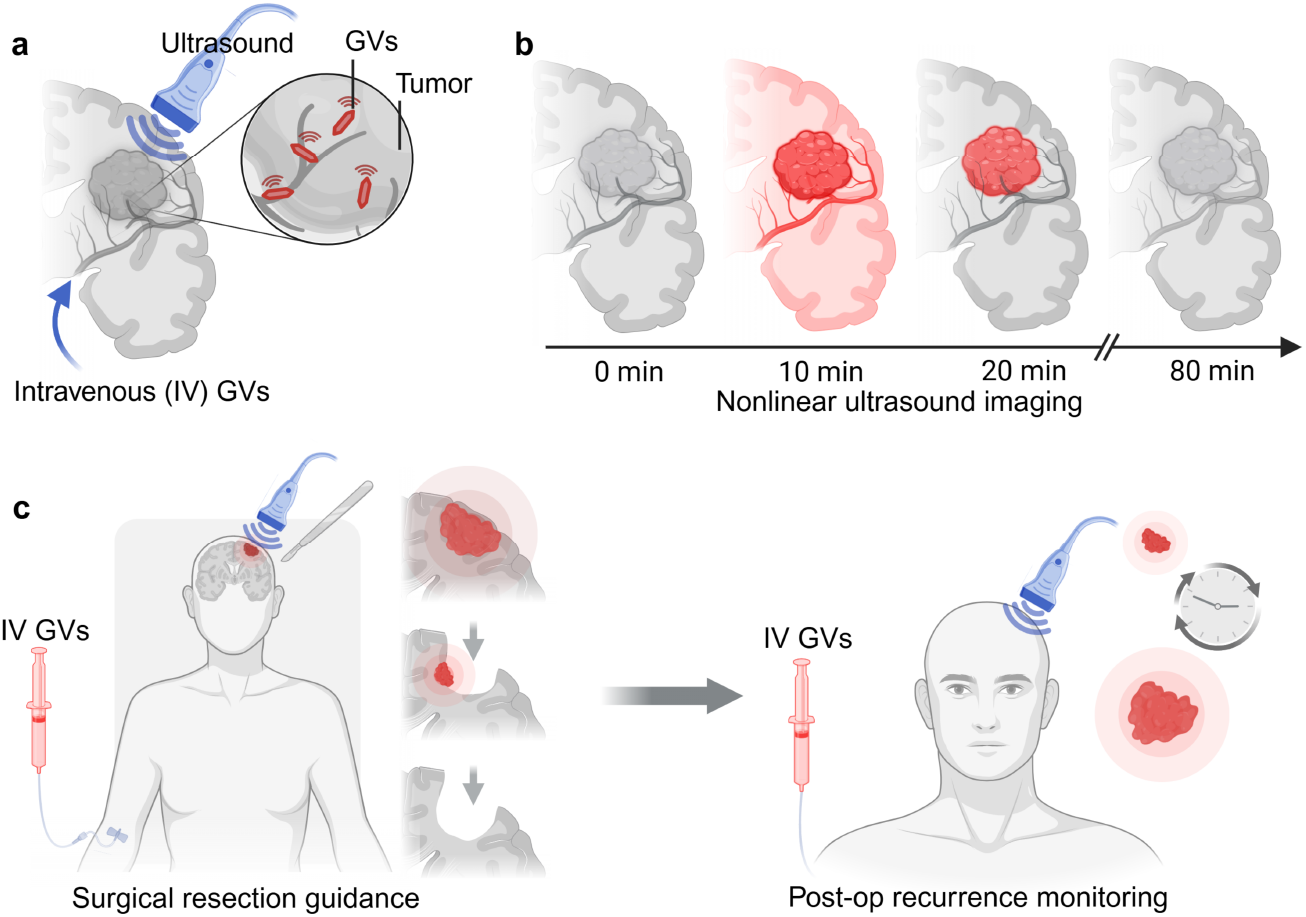
Envisioned use of gas vesicles as an acoustic tumor paint. **a**, Gas vesicles (GVs) are air-filled nanostructures that provide robust nonlinear ultrasound signal. When infused intravenously, they preferentially extravasate out of leaky tumor neovasculature into neoplastic tissue. **b**, Following GV injection, diffuse ultrasound signal is observed across the entirety of the imaging plane. GVs are then cleared from circulating blood by the liver, while signal remains within the tumor parenchyma. Eventually, the intratumoral GVs are degraded by local tumor-associated macrophages. **c**, Due to the preferential localization of GVs to brain tumors, this imaging paradigm can be used for intraoperative resection guidance and longitudinal postoperative recurrence monitoring.

## RESULTS

### GVs accumulate in brain tumors following IV injection

To assess whether GVs selectively accumulate in brain tumors, we administered purified nonlinear *Anabaena flos-aquae* GVs^46,47^ in saline into the tail vein of anesthetized C57BL/6 mice previously been implanted with syngeneic GL261 brain tumors. Using a nonlinear xAM pulse sequence^34^ to image through a craniotomy, we selectively visualized GV signals against tissue background and tracked their dynamic distribution within the brain (**Supplementary video 1**). A complementary power Doppler scan^48^ was also performed to visualize the vascular structure within the imaging volume, providing a spatial reference for GV accumulation. Upon injection, we observed that GV signals rapidly accumulated within the brain tumors, reaching easily detectable levels within approximately 10 minutes (**Fig. 2a**). The signal continued to increase in the tumor region, clearly delineating the tumor boundaries, suggesting that GVs were effectively extravasating into the tumor microenvironment. Over the next 80 minutes, the GV signal gradually declined to baseline levels, indicating the clearance or degradation of the GVs. In contrast, when we examined a region of equivalent size in the contralateral (non-tumor) hemisphere, the GV signal reached a peak intensity that was ninefold lower than in the tumor and returned to baseline within 20 minutes post-injection (**Fig. 2b**). The background GV signal can be attributed to GVs circulating in the bloodstream during the 5-minute infusion period, with subsequent clearance from circulation by the liver^30,49^. The stark difference in signal intensity and persistence between tumor and non-tumor regions highlights the selective accumulation of GVs in tumor tissue, enabling effective imaging while also highlighting the local biodegradability of this protein-based contrast agent.

**Figure 2:**
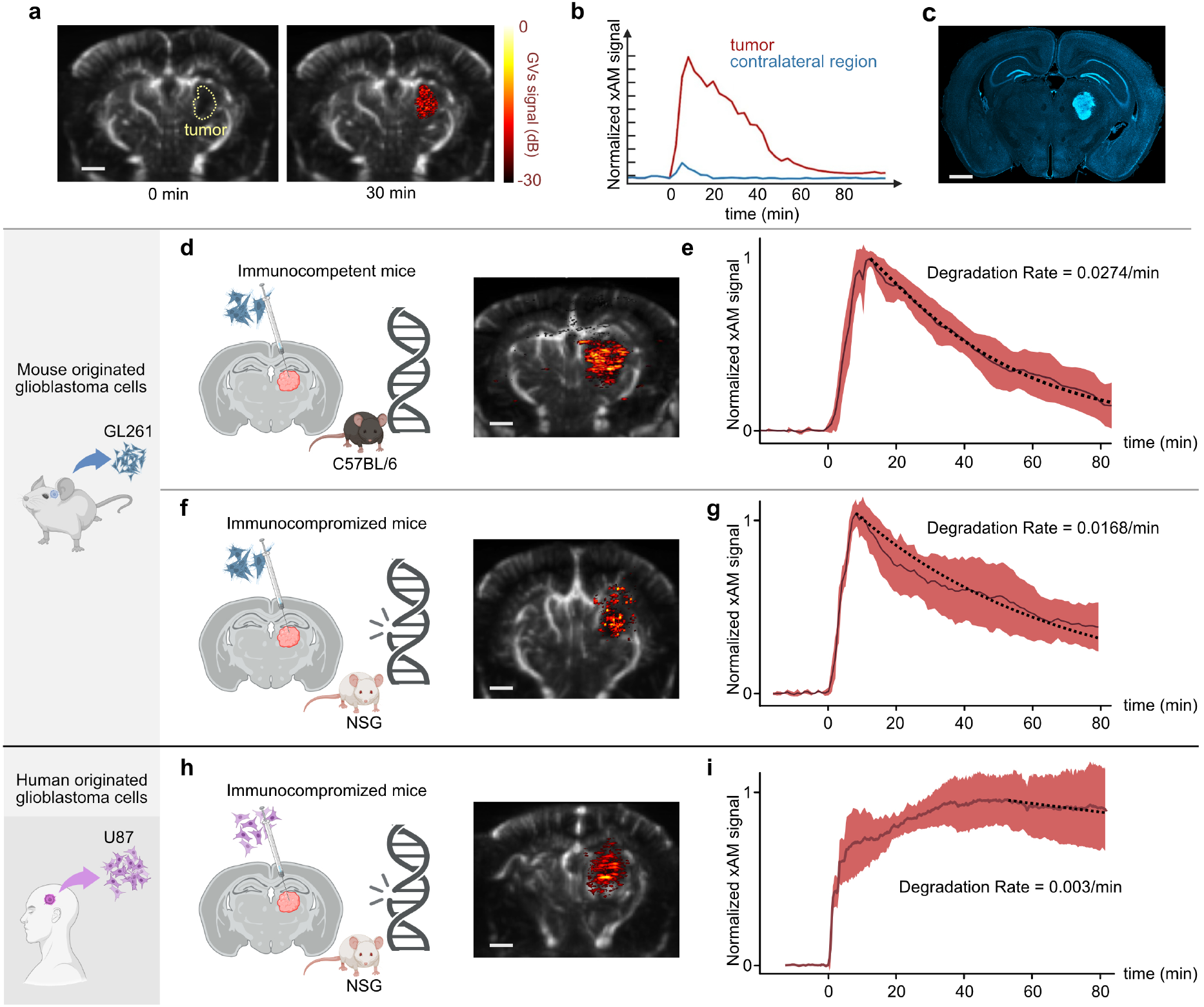
GVs selectively accumulate in brain tumors. **a**, Ultrasound images showing the overlay of brain vasculature (Doppler imaging, grayscale) and GV contrast (nonlinear signal, ‘hot’ colormap) in the coronal plane (Bregma −2m) of the brain of a representative mouse at day 7 post-tumor implantation. Following IV infusion GVs localize selectively to the tumor while being cleared from healthy vasculature. **b**, GV signal intensity within the tumor peaks approximately 10 minutes post-injection and gradually returns to baseline by 80 minutes. In contrast, the contralateral region exhibits a peak signal ninefold lower, returning to baseline within 20 minutes. **c**, Histological analysis of the corresponding brain slice (Bregma −2 mm) stained with DAPI demonstrates that the tumor morphology closely aligns with the GV-generated “acoustic painting.” **d, f, h**, Doppler and GV contrast overlays, alongside **e, g, i**, Signal kinetics, presented for three groups of mice: immunocompetent mice with GL261 tumors, immunocompromised mice with GL261 tumors, and immunocompromised mice with U87 tumors, all imaged at day 8 post-tumor implantation (N=5 per group). Solid lines indicate the mean time course averaged across mice in each group, with shaded areas representing ± standard deviation. Black dotted lines illustrate degradation kinetics modeled by the pharmacokinetic equation: dL/dT = k1. kc. B − k2. L where B is the blood signal, L is the liver signal, k1 is the uptake rate, k2 is the degradation rate, and kc is a signal scaling factor. Scale bar = 1mm.

Histological analysis confirmed the specificity of GV tumor labeling, showing that the brain area ‘painted’ by GVs corresponded precisely with the tumor region marked by cell nucleus (DAPI) staining (**Fig. 2c**).

In a separate control experiment, microbubbles, the standard ultrasound contrast agents used in clinical practice, were administered to mice with GL261 tumors, but no tumor-specific acoustic labeling was observed (**Supplementary Fig. 1**). This confirms that, unlike GVs, microbubbles are too large to extravasate out of vasculature into the tumor.

To evaluate the generalizability of GV-based tumor painting, we compared its performance across three different tumor models, including immunocompetent C57BL/6 mice with syngeneic GL261 tumors and immunocompromised NOD Scid Gamma (NSG) mice implanted with either syngeneic GL261 cells or xenografted human glioblastoma (U87) cells (**Fig. 2d, f, h**). In all three models, we consistently observed robust and selective GV accumulation within the tumor. Notably, NSG mice, which have defective macrophage function^50^, exhibited slower GV signal decay in the tumor compared to immunocompetent mice bearing the same GL261 tumors, with clearance rates of 0.0168 min^−1^ in NSG mice versus 0.0274 min^−1^ (**Fig. 2e, g**). Among the NSG group, mice with U87 tumors showed an even slower clearance rate (**Fig. 2i**).

These observations led us to hypothesize that the gradual reduction of GV signal in tumors is driven by degradation of GVs by tumor-associated macrophages (TAMs), analogous to their processing by Kupfer cells in the liver^30^. Among our three models, syngeneic GL261 tumors in immunocompetent mice are expected to have the highest density of TAMs. In contrast, NSG mice are deficient in functional macrophages, which would reduce GV clearance in both GL261 and U87 tumors. Additionally, U87 tumors, compared to GL261 tumors, are known to have lower TAM density ^51–53^. These findings support a role for TAMs in clearing GVs from the tumor microenvironment.

To more closely examine GV interactions with the tumor microenvironment, we collected brain tissues from immunocompetent GL261-implanted mice at multiple time points following infusion with GVs pre-labeled with an organic fluorophore, enabling detection through fluorescence microscopy (**Fig. 3**). Confocal fluorescence imaging of brain sections revealed a time-dependent increase in GV localization within the tumor. Quantitative mapping of GV spatial distribution in the histology slices revealed that 5 minutes post-injection, 34% of observed GVs were localized within the tumor, increasing to 73% at 25 minutes and 94% by 45 minutes (**Fig. 3a**). These histological findings closely matched the temporal dynamics of our ultrasound recordings.

**Figure 3:**
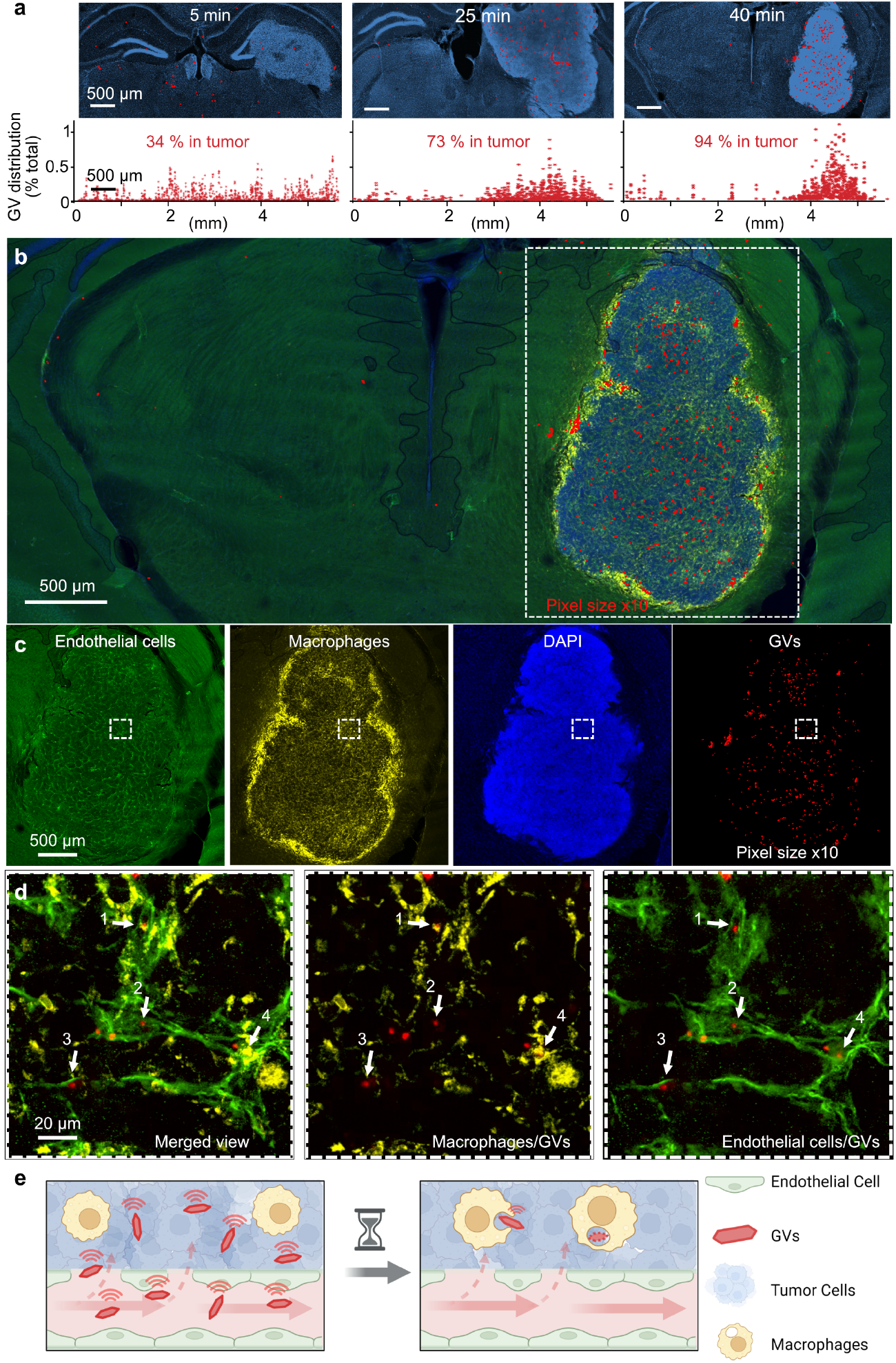
GVs localize in brain tumor parenchyma and are taken up by tumor-associated macrophages. **a**, (Top row) Histological images of GL261 tumors in mice at 5, 25, and 40 minutes post-injection of fluorescently labeled GVs. (Bottom row) Spatial distribution of GVs as a percentage of the total detected GVs along the horizontal axis of the corresponding brain slices shown above. **b, c**, Histological analysis of GL261 tumors in C57BL/6 mouse brains stained for endothelial cell marker CD31 (green), macrophage marker CD68 (yellow), DAPI (blue), and GVs (red), illustrating the localization of GVs within the tumor microenvironment. **d**, Magnified view of the boxed region in panel **c**, highlighting GVs in different degradation stages. Arrows 1 and 4 indicate GVs undergoing degradation by macrophages, while arrows 2 and 3 denote GVs sequestered in endothelial cells, not yet degraded by macrophages. **e**, Schematic representation of tumor-associated macrophage (TAM)-mediated GV degradation, explaining the observed signal kinetics.

To identify the specific sites of GV accumulation within tumor tissue, we performed immunostaining on brain sections collected 40 minutes post-GV injection using markers of endothelial cells (CD31) and macrophages (CD68) (**Fig. 3b**). In GL261 tumors, endothelial cells exhibited a characteristic “cracking” pattern indicative of angiogenic growth and BBB disruption^54^, a hallmark of aggressive tumors (**Fig. 3c**). Macrophages were prominent within the tumor core and at the tumor-healthy tissue interface, aligning with known patterns of TAM infiltration^55,56^. DAPI nuclear staining revealed the elevated cellular density typical of gliomas.

Within this microstructural environment, GVs were often found deep within the tumor mass, frequently positioned adjacent to disrupted blood vessels (arrows 2 and 3, **Fig. 3d**) and, in some cases, colocalized with macrophages (arrows 1 and 4, **Fig. 3d**). These observations suggest that GVs extravasate passively into the tumor through compromised neovasculature and are subsequently taken up and degraded by TAMs^57–59^.

### Acoustic tumor paint enables ultrasound-guided resection

After establishing the effective and generalizable labeling of brain tumors with GVs for nonlinear ultrasound imaging, we tested their use in ultrasound-guided neurosurgery. With current intraoperative neuronavigation, brain shift presents a persistent obstacle, sometimes hindering complete tumor resection; fluorescence-guided surgery using optical dyes has variable uptake depending on tumor type and is limited by light penetration ^9–15^. In contrast, ultrasound, already employed for anatomical guidance in neurosurgery, could help surgeons visualize tumors in real time if paired with an appropriate molecular tumor label. Critically, in this application, the contrast agent needs to persist during extended ultrasound exposure, a key property of GVs that is not typically shared by bubble-based agents^60^.

To evaluate the feasibility of neurosurgical guidance using acoustic tumor paint, we implanted mice with GFP-expressing U87 tumors and performed resections using surgical aspiration and sharp dissection after 9 to 12 days of tumor growth. Histological analysis of the *en bloc* resected tissue was performed to verify its tumor origin (**Fig. 4a**). Pre-resection ultrasound images clearly delineated GV-labeled tumors (**Fig. 4b**). After targeted aspiration and GV re-infusion, the absence of tumor labeling confirmed the successful removal of the tumor (**Fig. 4b**). This approach enables precise real-time visualization, even accounting for brain shift, significantly enhancing the accuracy and precision of tumor resection.

**Figure 4.**
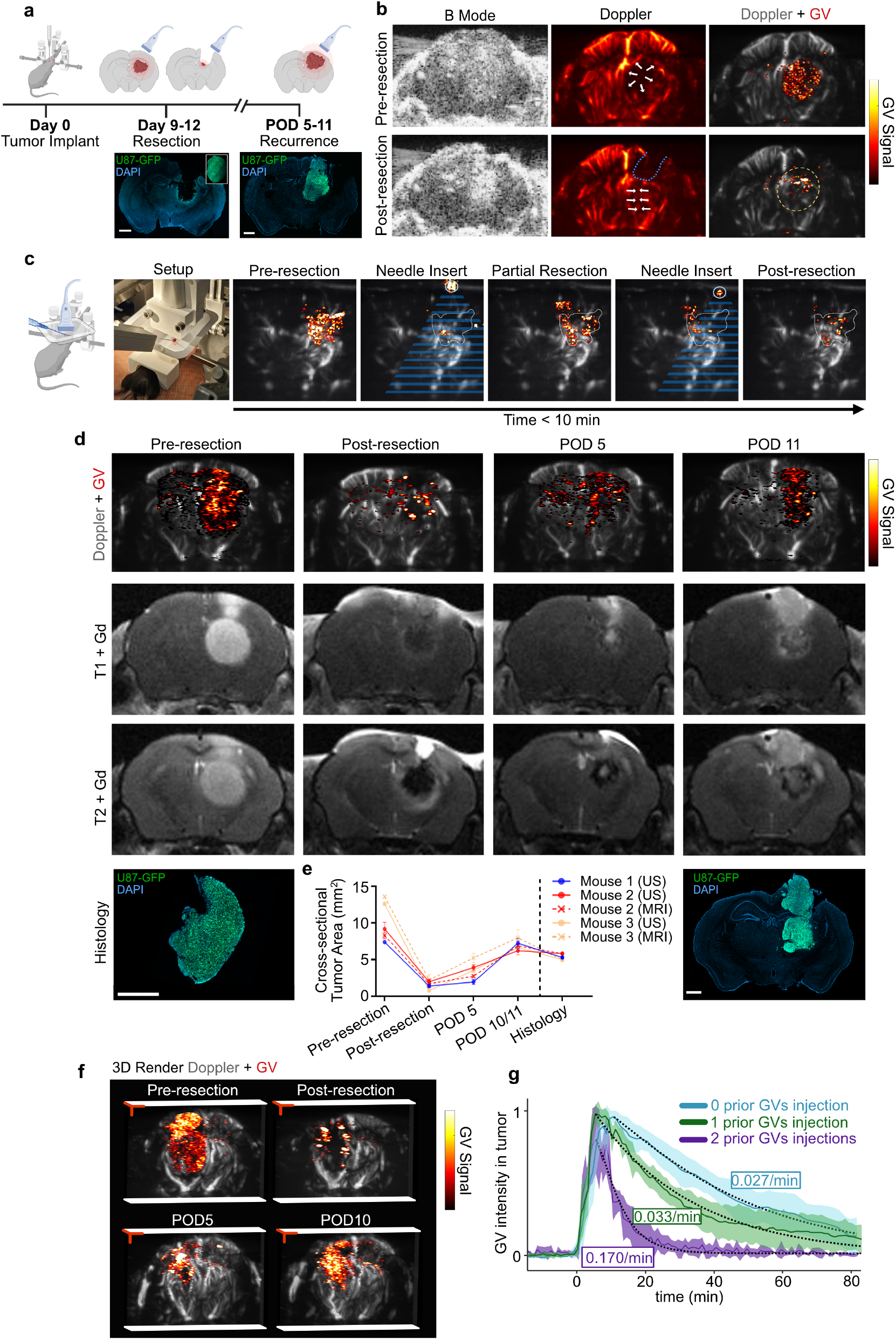
Acoustic tumor paint enables guided tumor resection and recurrence monitoring. **a**, Overview of tumor resection guidance and recurrence monitoring. Histological analysis of post-resection brain tissue, resected tumors, and recurrent tumors (in a separate mouse) demonstrates the effectiveness of the resection procedure and establishes a recurrence model. **b**, B-mode, power Doppler, and Doppler + GV nonlinear imaging before and after tumor resection. On B-mode imaging, diffuse hyperechoic signals are visible. Power Doppler reveals midline shifts and vasculature displacement due to the tumor’s mass effect (white arrows). Post-resection Doppler imaging shows only part of the resection cavity (blue outline) due to tissue recoil. GV nonlinear imaging distinctly delineates the tumor from healthy parenchyma on pre-resection images and highlights the extent of resection on post-resection images. **c**, Real-time ultrasound-guided tumor resection. Images captured pre-resection, during initial needle insertion, after partial tumor resection, during subsequent needle insertions, and post-resection. The needle (white circle) is echogenic, creating a shadowing effect (blue cone). **d**, Representative imaging of a mouse using Doppler + GV nonlinear imaging and gadolinium-enhanced T1- and T2-weighted MRI at pre-resection, post-resection, postoperative day (POD) 5, and POD 11. GV signal corresponds closely with T1-weighted MRI contrast. Histological analysis of the resected tumor on the day of surgery and recurrent tissue on POD 11 shows consistent tumor morphology. **e**, Cross-sectional tumor area (mean ± SEM) calculated for ultrasound (US) and MRI at all imaging time points, with histological measurements at the endpoint. **f**, 2D GV-Doppler overlay acquisitions were reconstructed into 3D representations of the tumor and vasculature at each time point for a representative mouse. **g**, Repeated GV administration effects on clearance dynamics. Clearance rates were faster in mice with one or two prior GV injections (0.033 min^−1^ and 0.170 min^−1^, respectively) compared to GV-naive mice (0.027 min^−1^). Scale bar = 1 mm.

To further assess the feasibility of using acoustic tumor paint for continuous procedural guidance, we conducted real-time imaging throughout the tumor resection process, capturing images before resection, during insertion of the vacuum-aspiration needle, throughout tumor debulking, and immediately following resection (**Fig. 4c**). The needle was clearly visible under nonlinear ultrasound imaging due to its strong specular reflection, allowing the monitoring of its position relative to the tumor. By the end of the resection, more than 90% of the tumor labeled by the GVs had disappeared (in signal intensity) from the area delineated during the pre-resection imaging (white dotted line). These findings demonstrate the potential of GV-based imaging to provide intraoperative guidance for tumor removal or biopsy procedures.

### Acoustic tumor paint enables relapse monitoring with ultrasound

Beyond providing real-time procedural guidance, ultrasound imaging combined with acoustic tumor paint could enable postoperative recurrence monitoring—an essential aspect of care for frequently recurring GBM tumors. Compared to established MRI protocols, ultrasound offers advantages such as greater accessibility, lower cost, and the convenience of point-of-care use. To explore the feasibility of this approach, we conducted longitudinal imaging of mice following tumor resection, assessed tumor detection limits, evaluated the health effects of repeated systemic GV injections, and tested GV imaging through human skulls and skull reconstruction materials.

We initiated longitudinal monitoring of mice before and immediately after tumor resection, continuing imaging on postoperative day (POD) 5 and either POD 10 or 11. To benchmark the performance of acoustic tumor paint against the clinical standard, we also conducted gadolinium-enhanced T1-weighted and T2-weighted MRI on the same animals. Imaging planes were aligned across sessions and with MRI using vascular structures (**Supplementary Fig. 2b**) and external fiducials. Our findings revealed a strong quantitative correspondence between the regions of ultrasound contrast seen with acoustic tumor paint and Gd-enhanced T1-weighted MRI (**Fig. 4d, e, Supplementary Fig. 2**).

Histological examination of the brain following recurrence confirmed that the tumor morphology and cross-sectional area matched both ultrasound and MRI images (**Fig. 4e**). To capture the full 3D extent of a recurrent tumor mass, we translated our linear array transducer along the elevation axis, acquiring a series of coronal images 0.2 mm apart. By compiling these sequential images, we generated a volumetric reconstruction of the tumor along with its surrounding vasculature (**Fig. 4f**).

In evaluating the translational potential of GV-based tumor painting, we assessed the systemic tolerability of repeated GV injection in both C57BL/6 and NSG mouse strains, assessing weight variation (**Supplementary Fig. 3A**), immunoglobulin levels (**Supplementary Fig. 3B**), and a metabolic panel (**Supplementary Fig. 3C**). No significant changes were observed in weight or blood biomarkers following single or repeated GV administration. Additionally, we examined the impact of repeated administration on tumor “painting” dynamics, given the production of anti-GV IgG and IgM antibodies (**Supplementary Fig. 3B**) ^27^. GVs were cleared more quickly in mice that had received one or two prior injections (clearance rates of 0.033 min^−1^ and 0.170 min^−1^, respectively) compared to GV-naive mice (clearance rate: 0.027 min^−1^) (**Fig. 4g**). While these accelerated kinetics are still compatible with recurrence monitoring, future modifications could reduce anti-GV antibody formation through surface passivation with endogenous proteins or inert polymers like polyethylene glycol^27,61–63^. Moreover, it is not uncommon for clinically-approved US contrast agents, such as microbubbles, to undergo accelerated blood clearance due to anti-PEG IgM and IgG production after repeated administrations^64^.

Assessing the influence of tumor size on tumor paint effectiveness, relevant to detecting recurrence or regional metastasis, we imaged immunocompetent mice at 3, 7, and 15 days after syngeneic tumor implantation (**Fig. 5a**). Distinct GV accumulation was detectable in brain tumors as early as the 3-day time point, when tumors were approximately 1 mm in diameter (based on histology) and ultrasound-identified tumor location was confirmed by histology. By day 7, tumors had grown to around 2 mm and displayed uniform GV accumulation. By day 15, tumor size frequently exceeded 5 mm, and GVs accumulated primarily in viable, vascularized, non-necrotic tumor regions. Kinetic analysis revealed that GV signal decayed most rapidly in 3-day-old tumors (clearance rate: 0.051min^-1^), with progressively slower rates in 7-day-old tumors (clearance rate: 0.025 min^-1^) and 15-day-old tumors (clearance rate: 0.006 min^-1^) (**Fig. 5b**). This may reflect heightened macrophage activity in early tumor growth stages. Across these variations, tumor paint effectively detected and delineated tumors as small as 1 mm, up to the maximum size compatible with the mouse model, demonstrating its utility in postoperative, longitudinal recurrence monitoring comparable to more expensive MRI scans.

**Figure 5:**
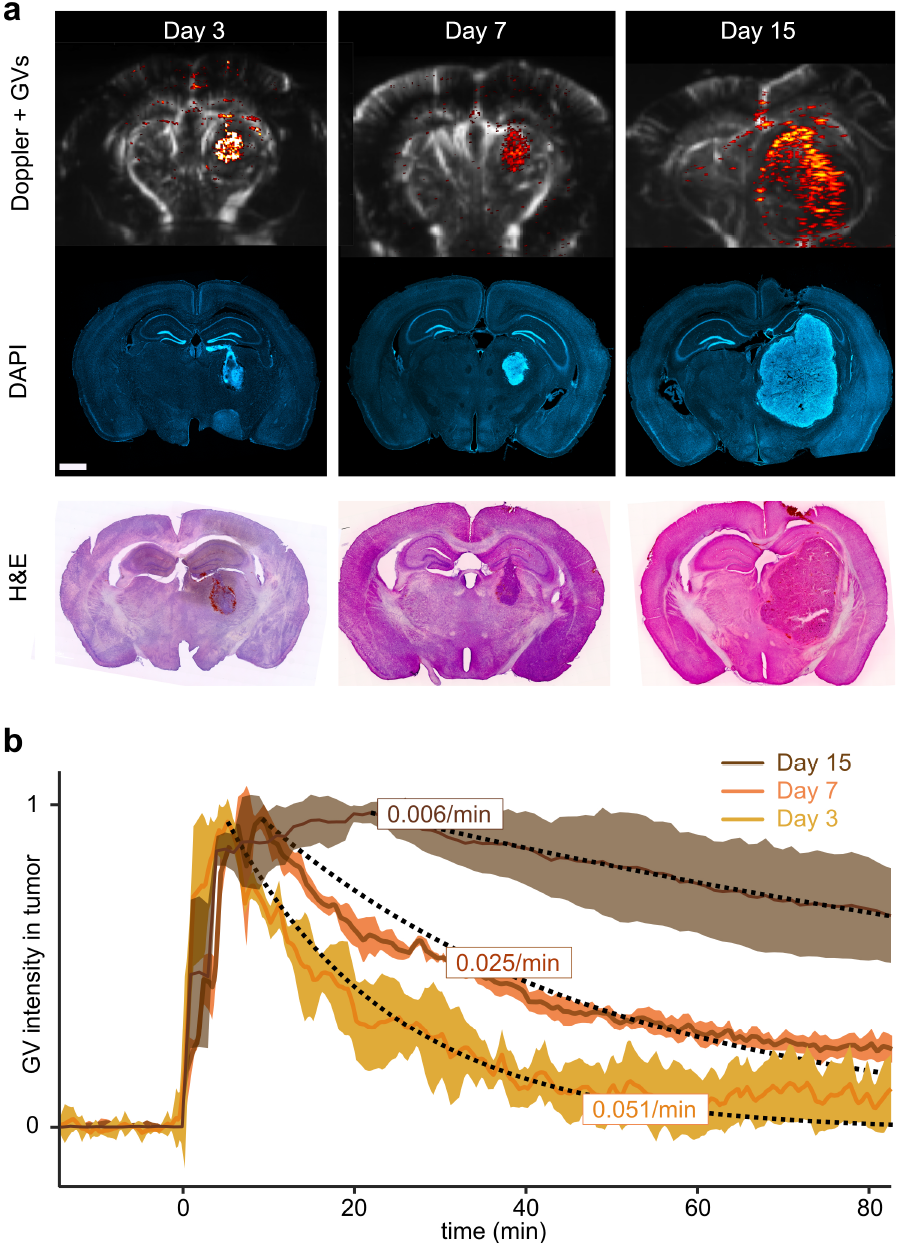
Acoustic tumor paint can detect tumors as small as 1 mm. **a**, Representative ultrasound images and corresponding histological staining (DAPI and hematoxylin and eosin) demonstrating GV accumulation in brain tumors at days 3, 7, and 15 post-GL261 tumor implantation in mice. At day 3, GV accumulation is detectable in small tumors (∼1 mm) and confirmed by histology. By day 7, tumors (∼3 mm) display uniform GV distribution, while at day 15, larger tumors (<5 mm) exhibit heterogeneous GV distribution, with minimal GV presence in necrotic regions. **b**, Kinetic analysis of GV clearance at different stages shows the fastest clearance in 3-day-old tumors (0.051/min), a moderate rate in 7-day-old tumors (0.025/min), and the slowest clearance in 15-day-old tumors (0.006/min). N=4 mice/group. Scale bar = 1 mm.

### Feasibility of GV-based imaging through human skull and cranial prosthesis

While intraoperative imaging does not require penetration of the skull, effective postoperative recurrence monitoring relies on the ability to detect GV signals through either an intact skull or a cranial prosthesis, which may be installed during initial tumor resection (**Fig. 6a**). To evaluate this capability, we imaged GVs through both a cadaveric human skull and an FDA-approved cranial implant made of polymethyl methacrylate (PMMA) (**Fig. 6b**). PMMA is widely used in cranial implants due to its biocompatibility, mechanical durability, and superior acoustic transparency compared to natural bone, allowing for high-resolution transcranial imaging^65–69^. This property makes PMMA an ideal candidate for facilitating GV-based imaging in clinical settings^69^.

**Figure 6:**
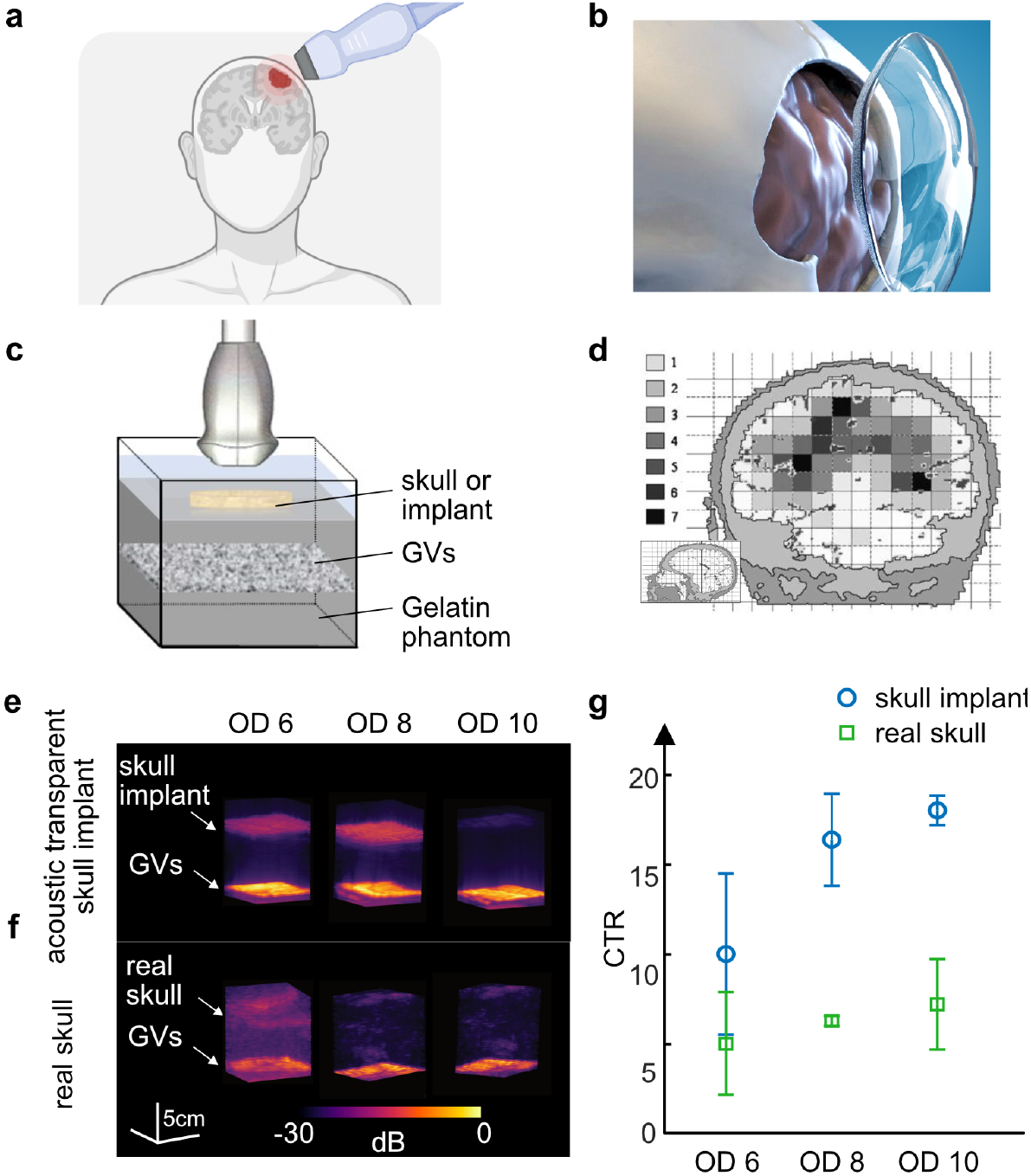
GVs can be detected with ultrasound through human skull and sonolucent cranial prosthesis. **a**, Schematic illustration of transcranial ultrasound diagnosis for brain tumors. **b**, Acoustic transparent cranial implant fabricated from PMMA (adapted from longeviti.com). **c**, Experimental setup showing a matrix probe recording signals from a tissue-mimicking phantom embedded with GVs, positioned beneath either the acoustic transparent cranial implant or a real human skull. **d**, Coronal projection of glioma distribution within the brain, showing the anatomical site distribution. Colors indicate the number of gliomas in each 1×1-cm square, with smoothing applied across adjacent squares. The inset depicts the section plane (adapted from Larjavaara et al., Neuro Oncol, 2007). **e, f**, Ultrasound images of the GV-embedded phantom captured through the acoustic transparent cranial implant (e) and through a real human skull (f). **g**, Contrast-to-noise ratio (CTR) quantification of the ultrasound data.

For transcranial imaging experiments, we employed a lower-frequency matrix array probe (1.25 MHz) to minimize attenuation, achieve greater depth, and capture volumetric images. The probe was positioned above either a human skull segment or the PMMA skull implant (**Fig. 6c**). Both were placed atop a tissue-mimicking phantom containing a 5 mm-thick layer of GVs at 6 cm depth below the skull or implant, approximating typical GBM tumor locations and dimensions at initial diagnosis or recurrence^52,53,70^ (**Fig. 6d**). For the PMMA implant, this setup models a clinical scenario in which the implant is installed post-resection, allowing subsequent imaging to monitor for recurrence. The GV concentrations within the phantom inclusions matched those evaluated in tumors (**Supplementary Fig. 4**). Our results demonstrated clear GV contrast through both the human skull and PMMA implant (**Fig. 6e, f**). As expected, contrast was higher with the PMMA implant, but even through the human skull, and at the lowest GV concentration tested, the contrast-to-tissue ratio (CTR) remained above 5. These findings support the feasibility of acoustic tumor paint for post-surgical monitoring and suggest that replacing the removed skull segment with an acoustically transparent material like PMMA may offer significant advantages for ongoing surveillance of tumor recurrence.

## DISCUSSION AND CONCLUSION

In this study, we introduced and validated acoustic tumor paint, a novel ultrasound imaging approach that leverages the high intratumoral accumulation of GVs to enhance the diagnosis, surgical resection, and recurrence monitoring of GBM tumors. Our findings confirm the fundamental feasibility of this approach, elucidate its mechanisms, demonstrate its performance across multiple GBM mouse models, and establish robust cross-validation with gold-standard MRI and histology methods. We also demonstrated its safe, repeatable use, and detectability across human skull and prosthetic materials, supporting its potential for clinical applications. The ability to provide high-resolution intraoperative imaging would offer neurosurgeons a real-time view of tumor localization, while the capacity for acoustic recurrence monitoring would provide a more accessible alternative to MRI.

A key strength of acoustic tumor paint is the tumor-specific accumulation of GVs in the tumor microenvironment following intravenous injection, followed by degradation by TAMs (along with possibly other resident cells such as glia, which should be further investigated). The rapid uptake, sustained retention, and compatibility with continuous non-destructive nonlinear imaging together define an optimal imaging window for diagnostic monitoring and real-time procedure guidance. Furthermore, this approach could be integrated with functional ultrasound imaging during surgery, allowing for real-time hemodynamic mapping to help preserve critical functions following resection^69^. As a further surgical aide, fluorescent labeling of GVs could enable optical identification of tumor tissue in the resection cavity.

GVs also show promise for longitudinal monitoring due to their compatibility with repeated systemic administration, which is particularly advantageous over MRI for patients in resource-limited environments or those with severe health constraints. If this approach proves valuable in clinical settings, we anticipate that the use of sonolucent skull prostheses may become more widespread following tumor resection, providing superior resolution and sensitivity in post-surgical monitoring.

To advance clinical translation, it will be essential to establish clinical-grade production of GVs and validate their safety in larger animal models, drawing from the extensive experience with other protein-based diagnostics and therapeutics. Reducing or delaying anti-GV immune responses will also be important for long-term monitoring, which could be achieved by functionalizing the GV surface through genetic or chemical modification to alter the protein corona. Future work may also explore functionalizing GVs for enhanced specificity — such as targeting tumor-specific antigens or sensing molecular markers in the tumor microenvironment — to increase tumor accumulation and imaging specificity even further.

With these improvements and further pre-clinical and clinical validation, we can picture acoustic tumor paint substantially improving surgical outcomes and assisting tumor management from diagnosis to long-term monitoring.

## MATERIALS AND METHODS

### Gas vesicle preparation as an injectable contrast agent

Nonlinear GVs were prepared as previously described^47^. Briefly, GVs were harvested from buoyant *Anabaena flos-aquae* cells by hypertonic lysis and purified by repeated centrifugally-assisted flotation and resuspension. The GVs were stripped of their outer GvpC layer by treatment with 6M urea, followed by additional repeated centrifugally-assisted flotation, dialysis in 1x phosphate buffered saline (PBS), and resuspension to remove the GvpC and urea. This enables buckling of the GV shell resulting in a nonlinear ultrasound signal. For fluorescent labeling, the purified GVs were incubated with Alexa Fluor™ 594 NHS Ester at a concentration of 800 x molar excess for 2 hours at room temperature. The labeling reaction was quenched by adding 1M Tris buffer at a 1:100 volume ratio to the labeling mixture. Then, the labeled GVs were dialyzed in 1 x PBS for four rounds in order to remove excess Alexa Fluor™ 594 NHS Ester (Succinimidyl Ester).

### Overview of mouse experiments and surgery

All procedures were approved by the Institutional Animal Care and Use Committee of the California Institute of Technology. Adult NSG or C57BL/6 female mice aged 6-14 weeks (Jackson Laboratory) were used; no randomization or blinding was necessary for this study. The mice were anesthetized with 4% isoflurane with room air for induction, followed by maintenance at 1-2% isoflurane using a stereotaxic instrument. Core body temperature was maintained at 37°C using a heating pad. Local analgesics (0.25% bupivacaine, 1 mg/kg, subcutaneous) and systemic analgesics (ketoprofen, 5 mg/kg; buprenorphine, 1 mg/kg, both subcutaneous) were administered preoperatively. Following surgery, the mice were monitored and remained bright, alert, and responsive throughout the postoperative period. Cage water was supplemented with antibiotics (trimethoprim/sulfamethoxazole) and analgesics (ibuprofen) for a week.

### Intracranial implantation of brain tumors for tumor paint characterization

Intracranial injections of GL261 and U87 tumor cells were conducted in immunocompetent c57bl/6 (for GL261 and U87 tumors) and immunocompromised NSG mice (for U87 tumors). GL261 and U87 cells were recovered from LN2 storage and cultured without drug selection for at least 2 passages before implantation. On the day of implantation, cells were dissociated using Trypsin/EDTA (Corning 25-053-CI), centrifuged at 500 g for 5 minutes, and resuspended in culture media at a concentration of around 60 million cells per milliliter. The cells were then stored on ice before injection.

After making a small burr hole using a micro drill steel burr (Burr number 19007-07, Fine Science Tools), the cells were injected intracranially at the coordinate of Anterior-Posterior (AP) −2 mm, Media-Lateral (ML) +1.5 mm, and Dorsal-Ventral (DV) −3.5 mm, relative to bregma. The injection was conducted using a microliter syringe (Hamilton) with a 33G needle (World Precision Instrument) controlled by a micro syringe pump (World Precision Instrument) at a flow rate of 5 - 7 nL per second. A volume varying between 2.5 to 5 nL containing 10^5^ cells was injected. The burr hole was covered with bone wax (Lukens), and the skin was closed after the injection with a tissue adhesive (GLUture).

### Intracranial implantation of brain tumors for surgical resection

We sought to establish a reliable tumor resection and recurrence model. On the day of tumor implant, a craniotomy was first performed from AP + 0.0 to AP - 3.5 mm and from ML - 2.5 to ML + 2.5 mm using a micro drill steel burr. Care was taken to not damage the dura. GFP-expressing U87 cells (2 × 10^5^ cells) were then implanted at AP - 1.4 mm, ML + 1.5 mm, and DV - 3.5 mm to ensure growth away from major blood vasculature and to enable ease of access during tumor resection. The skull flap was replaced with a sonotransparent TPX (polymethyl pentane) window of similar size and affixed in place using a UV-curable composite (Tetric EvoFlow) and dental metabond cement (Patterson Dental Company).

### Ultrasound imaging

Imaging was performed using a Verasonics Vantage programmable ultrasound scanning system and a L22-14vX 128-element linear array Verasonics transducer, with a specified pitch of 0.1 mm, an elevation focus of 8 mm, an elevation aperture of 1.5 mm and a center frequency of 18.5 MHz with 67% −6 dB bandwidth. For nonlinear image acquisition, a custom cross-amplitude modulation (xAM) sequence^34^ was used with an xAM angle (θ) of 19.5°, an aperture of 65 elements, and a transmitting frequency at 15.625 MHz. GVs were intravenously injected through a catheter in the lateral tail vein previously sterilized with ethanol and flushed with heparinized saline. Residual GVs in the catheter were also flushed with heparin to ensure delivery of correct volume.

The power Doppler images mapping cerebral blood volume (CBV) were acquired at 15.625 MHz. Briefly, the pulse sequence contains 15 tilted plane waves varying from −14° to 14° at a 500 Hz pulse repetition frequency. A block of 200 coherently compounded frames was processed using an SVD clutter filter to separate tissue signal from blood signal (cutoff of 40) to obtain a final power Doppler image exhibiting CBV in the whole imaging plane^48^.

### Imaging session for tumor paint characterization

Unless mentioned otherwise, the ultrasound imaging was conducted 8 days after the surgery. A cranial window from approximately AP - 1 mm to AP - 3 mm and from ML - 2.5 mm to ML + 2.5 mm was performed using a micro drill steel burr. Care was taken to not damage the dura.

The ultrasound transducer was held by a custom holder mounted on the arm of the stereotaxic instrument, and ultrasound coupling gel (Aquasonic) was directly applied to the cranial window. The transducer surface was placed ∼1 mm above the middle of the brain in the coupling gel. First, a power Doppler scan of the cranial window was performed along the anterior-posterior direction. Similarly, an xAM scan was applied at 426 kPa to scan across the cranial window before any GV administration. After acquiring this baseline, 1.3 pmol of purified stripped Anabaena flos-aquae GVs were then intravenously administered in a volume of 150 µl saline at a rate of ∼0.8µl/s, while continuously scanning the brain with xAM imaging at an imaging rate of ∼1 image/min per coronal plane. After ∼15 min, the coronal plane displaying the highest signal in the tumor was chosen for longitudinal imaging (total 100 min of imaging) with the imaging rate of ∼1 image/min.

### MRI Imaging

After anesthesia, the mice were placed in the custom-made plastic head mount and imaged in a 7 T MRI (Bruker Biospec). Subsequently, the mice were injected via the tail vein catheter with 0.125 µmol of ProHance (Bracco Imaging) dissolved in sterile saline, per gram of body weight. A T_1_-weighted rapid acquisition with refocusing echoes sequence (RARE) (echo time TE = 7.00 ms, repetition time TR = 750 ms) was used to verify and record the coronal tumor anatomy in coordination of the mouse brain. Lastly, a coronal T_2_-weighted RARE sequence (echo time TE = 33 ms, repetition time TR = 2500 ms) was also applied for implanted tumor lesion characterization as well as edema detection. Slice thicknesses were 0.2 mm for both MRI sequences.

### Imaging for tumor resection guidance and post-operative recurrence monitoring

Imaging was conducted after 9 - 12 days of tumor growth. Given significant blooming artifacts from the metal oxide in the metabond cement on MRI, the TPX window was removed and replaced for all resection and imaging sessions. On the day of resection, mice were anesthetized and maintained as described previously, and the TPX window was removed. Preoperative scans of the tumor were conducted with MRI and US. Using gentle vacuum aspiration and sharp dissection, the tumor was resected and hemostasis was achieved using an absorbable gelatin hemostatic sponge (Goodwill AGS111). After thoroughly irrigating the resection cavity with cold saline, another IV injection of GVs was administered and the same planes were imaged for remnant tumor (with both US and MRI). The TPX window was then replaced, and the mouse was allowed to recover. On POD 5 and either 10 or 11, the tumor was similarly imaged with both US and MRI following TPX window removal.

For all images in this section, TPX removal results in higher US signal from the cortex than traditionally seen, likely due to acute local inflammation; the skull base also had some nonlinear artifact. As a result, all nonlinear US images have identical masks within a given imaging day applied to remove ∼0.5 mm of cortex (**Supplementary Fig. 2a**). For computing the cross-sectional area of the tumor, the same mask was applied to US and MRI acquisitions and histology; segmentation and measurement was computed by n = 3 individuals and averaged. Between imaging sessions, planes were matched using vasculature, anatomical structures, and general tumor morphology (**Supplementary Fig. 2b**). A series of coronal images were taken 0.2 mm intervals apart and spanned the entire cranial window at each timepoint. We generated a 3D reconstruction of the tumor along with its surrounding vasculature using the python library Napari (DOI 10.5281/zenodo.3555620).

### Real-time imaging and resection

For simultaneous ultrasound imaging and tumor resection, a GL261 cortical tumor was implanted in a C57bl/6 mouse at AP - 1 mm, ML + 1.5 mm, and DV - 1 mm for ease of access. A syngeneic model was used for more effective cortical growth. After 14 days, a craniotomy was performed as discussed, and a small weight boat with a comparable-sized opening was cemented onto the skull and filled with PBS. The transducer was positioned as normal using one of the stereotax arms while a second stereotax arm was used to position the vacuum aspiration needle at around a 45° angle. Following GV injection, continual xAM imaging was conducted while the vacuum aspiration needle was advanced and retracted several times (to ensure proper debulking of tumor).

### Repeated GV injection for toxicity study

To assess the impact of repeated GV administration on GV degradation dynamics, we conducted a study with three groups, each consisting of N=4 mice.

#### Control group (0 prior GV injections)

Mice in this group were intravenously injected with 1.3 pmol of purified, stripped *Anabaena flos-aquae* GVs in 150 µl of saline on day 0 and imaged immediately afterward following the protocol described previously.

#### 1 prior GV injection group

Mice received the same quantity of GVs on day 0 and day 7. Imaging was performed immediately after the second GV injection.

#### 2 prior GV injections group

Mice were administered the same quantity of GVs on days 0, 7, and 14, with imaging conducted immediately after the third GV injection.

### Collection of brain tissue

After the imaging session, animals were perfused with 30 mL of PBS, followed by 30 mL of 10% formalin solution. The brains were resected and placed in 10% formalin solution for 24-36 hours at 4°C after which they were transferred to PBS for long-term storage at 4°C. For frozen sectioning, samples were immersed in 15% and 30% sucrose/PBS solution for 12 - 24 hrs each and were then embedded in OCT and frozen overnight at −80°C before sectioning.

### Histology of implanted cells in mouse brain

Brain tissues were sectioned into 60 - 80 µm slices using a vibrating microtome (Leica) or a cryotome (Leica). The slices were stained with DAPI nuclear stain (Thermo Scientific 62248) or directly mounted with ProLong Diamond antifade mountant with DAPI (Thermo Fisher P36962). For immunofluorescent staining, the samples were first incubated with antigen retrieval solution, followed by blocking buffer and primary antibodies CD31 (Abcam AB56299) and CD68 (BioLegend 137001), relevant secondary antibodies, and mounted with a cover slip. Some sections were stained using hematoxylin and eosin (Abcam AB245880) and mounted with a limonene mounting medium (Abcam AB104141). The mounted slices were imaged using a Zeiss LSM 800/980 confocal microscopes with ZEN Blue. **Fig. 3b-c** images show maximum intensity projections of z-stack images (8 planes, Δz = 4 μm). **Fig. 3d** images display 3D volumes of z-stack images (13 planes, Δz = 3 μm), which better capture the structure of blood vessels compared to single z-plane images. Images were processed and exported using the ZEN Blue software.

### Volumetric transcranial imaging in GV phantoms

To determine the capabilities to image GVs volumetrically and transcranially, a gelatin-based ultrasound phantom was created. A 5 mm-thick layer of gas vesicles, suspended in 4% gelatin and 1X PBS mixture, was embedded at 6 cm depth in the larger phantom. The surrounding phantom consisted of 5% gelatin in deionized water. Three different concentrations of GVs were used – OD 06, OD 08, and OD 10 – with three phantoms created for each concentration. To image the phantoms, a 1.25 MHz sparse matrix array designed for volumetric transcranial human imaging with a 65 mm imaging aperture was used^71^. This transducer was controlled by a 1024-channel Verasonics volumetric vantage system. Datasets were acquired using a focused grid approach, where the sound was focused in a 20 × 20 grid at 65 mm in depth, with each focal point 0.6 mm from the previous, resulting in a 12 mm × 12 mm square. Each phantom was imaged through a piece of human temporal bone, placed 1 cm below the transducer, and a PMMA-based acoustically transparent skull implant (ClearFit®, Longevity, USA), placed 3 cm below the transducer surface. The positioning varied due to the smaller size of the implant and the need to keep it within the path of the ultrasound. Datasets were beamformed using delay and sum beamforming, with comparisons between concentration groups measured using the CNR. CNR was calculated as:

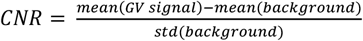

### Serum collection, IgG and IgM quantification, and complete metabolic panel

Both NSG and C57BL/6 mice were injected with either 150 uL of PBS or GVs (as described before). Blood was collected in Vacutainer™ Venous Blood Collection Tubes (BD) via a submandibular vein puncture on day 5 before PBS or GV injection and on days 2, 9, and 16 following PBS or GV injection. Serum was frozen down until ready to analyze. ELISA kits were used to quantify bulk IgG (Cayman Chemical 501240) and IgM (Abcam AB215085) levels. Samples were run in triplicates (mean + SEM) and diluted to stay within the range of the standard provided; appropriate controls were also run. Absorbance was quantified at 450 nm using a microplate reader (Tecan Spark).

Whole blood collection occurred on day 20 following GV injection, at which point the mouse was euthanized. Serum alanine aminotransferase (ALT), aspartate aminotransferase (AST), alkaline phosphatase, blood urea nitrogen, creatinine, total bilirubin, and total protein were quantified by VRL Animal Health Diagnostics.

### Microbubble imaging

GL261 tumors were implanted in c56bl/6 mice and imaged as previously described. At a concentration of 1.2 × 10^10^ bubbles per mL and with each bubble at 10.3 μm^3^ of gas, 150 ul of microbubbles (Definity)^72^ were injected intravenously and imaged using xAM to match the volume of gas injected via GVs.

## ACKNOWLEDGEMENTS

The authors would like to thank Di Wu, PhD for providing GFP-expressing U87 cells, Margaret Swift for her help with animal procedures and Linda Liau, MD, PhD, MBA and Chirag Patil, MD for helpful discussions. Diagrams were generated using BioRender.

## FUNDING

National Institutes of Health (RO1-EB018975 to MGS, RF1NS113285 to GFP).

Sontag Foundation: CR, GD, MGS

UCLA-Caltech MSTP (NIGMS T32 GM152342): GD

Human Frontier Science Program Cross-Disciplinary Fellowship (LT000217/2020-C): CR

M.G.S. is an investigator of the Howard Hughes Medical Institute.

## AUTHOR CONTRIBUTIONS

*Conceptualization:* All authors

*US sequence development:* CR, RVB, RJ, GP

*Rodent surgery and experiments:* CR, GD, HRL

*Histology and imaging:* CR, GD, PBL

*Toxicology and blood analysis:* CR, GD, PBL

*MR imaging:* GD, HRL

*In-vitro phantom experiments:* RVB, RJ, DM, GFP

*Writing - original draft:* CR, GD, MGS

*Writing - review & editing:* All authors

## DATA AND MATERIALS AVAILABILITY

Key data from this paper will be made available on CaltechDATA (https://data.caltech.edu/) or similar data repositories. DOI accession number will be generated upon acceptance of paper. Code used to generate key figures in the result section will be made accessible on a GitHub repository.

## SUPPLEMENTARY MATERIAL

**Video S1: Visualization of intravenously administered GVs in brain tumors**. Real-time tracking of GV circulation (nonlinear signal displayed with a ‘hot’ colormap) in the coronal plane (Bregma −2 mm) of a representative mouse at 7 days post-tumor implantation. Following intravenous infusion, GVs selectively accumulate in the tumor while clearing from healthy vasculature. Within the tumor, GV signal intensity peaks approximately 10 minutes after injection and gradually diminishes to baseline by 80 minutes. In contrast, the contralateral region shows a ninefold lower peak signal, normalizing within 20 minutes.

**Figure S1:**
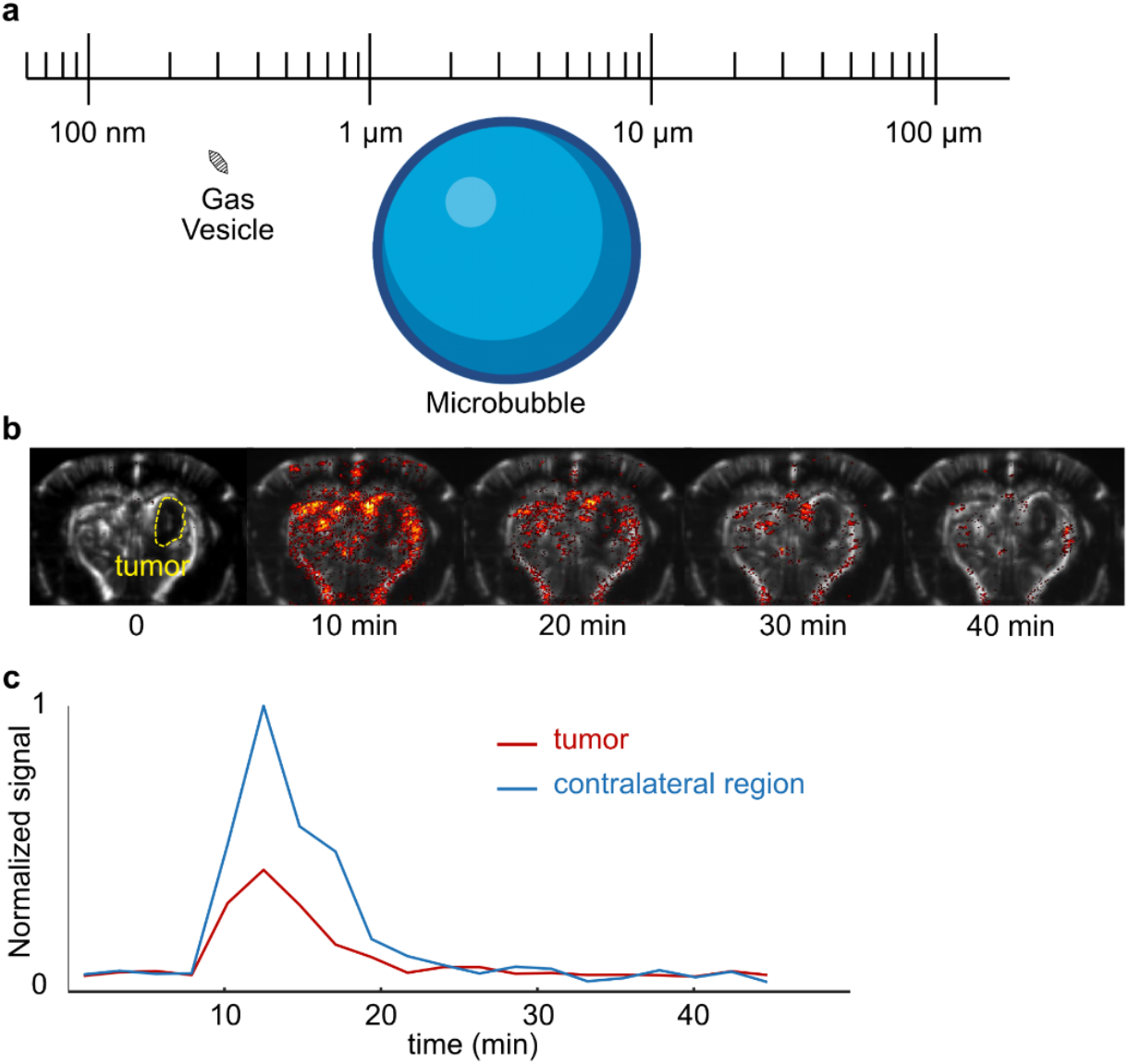
Microbubbles do not provide effective US tumor contrast. **a**, Microbubbles (ranging from 1-10 µm) are significantly larger than gas vesicles (around 85 nm by 500 nm) and are far too big to extravasate out of the tumor neovasculature. **b**, Tumor contrast enhancement imaging with intravenous microbubbles over time. There is minimal signal from the tumor core (outlined) at various time points. **c**, Signal intensity over time within the tumor compared to the contralateral side demonstrates the limited tumor-specific contrast.

**Figure S2:**
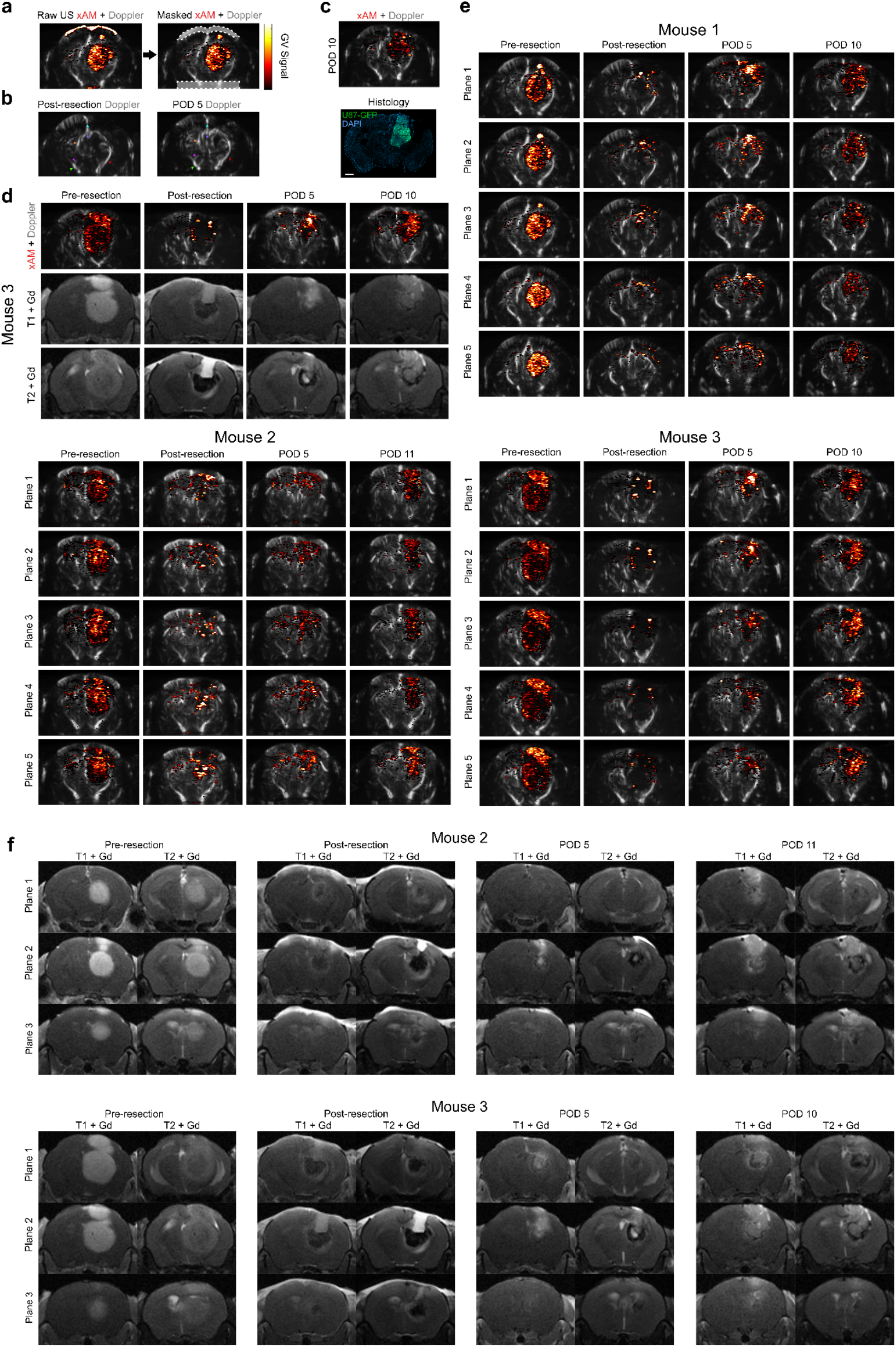
Additional representative mice demonstrating GV-imaging for tumor resection guidance and recurrence monitoring. **a**, A mask is applied to the nonlinear signal to remove artifacts on the surface of the brain resulting from TPX removal and the ventral aspect of the brain resulting from reflection from the skull base. **b**, In between imaging sessions, the same plane was reliably found by looking for features in doppler imaging. This is an example of the same mouse five days apart (post-resection and on POD5) with similarly-colored arrows indicating the same vascular structures. **c**, An additional mouse demonstrating similar recurrent tumor morphology on US and histology on POD 10. **d**, An additional mouse demonstrating close synergy between US and T1 / T2-Gd enhanced MRI. **e**, Anterior (Plane 1) to posterior (Plane 5) coronal US scans with step size of 0.2 mm in three representative mice. **f**, Contrast-enhanced MRI for mouse 2 and 3 across three planes at each of the timepoints. Scale bar = 1 mm.

**Figure S3:**
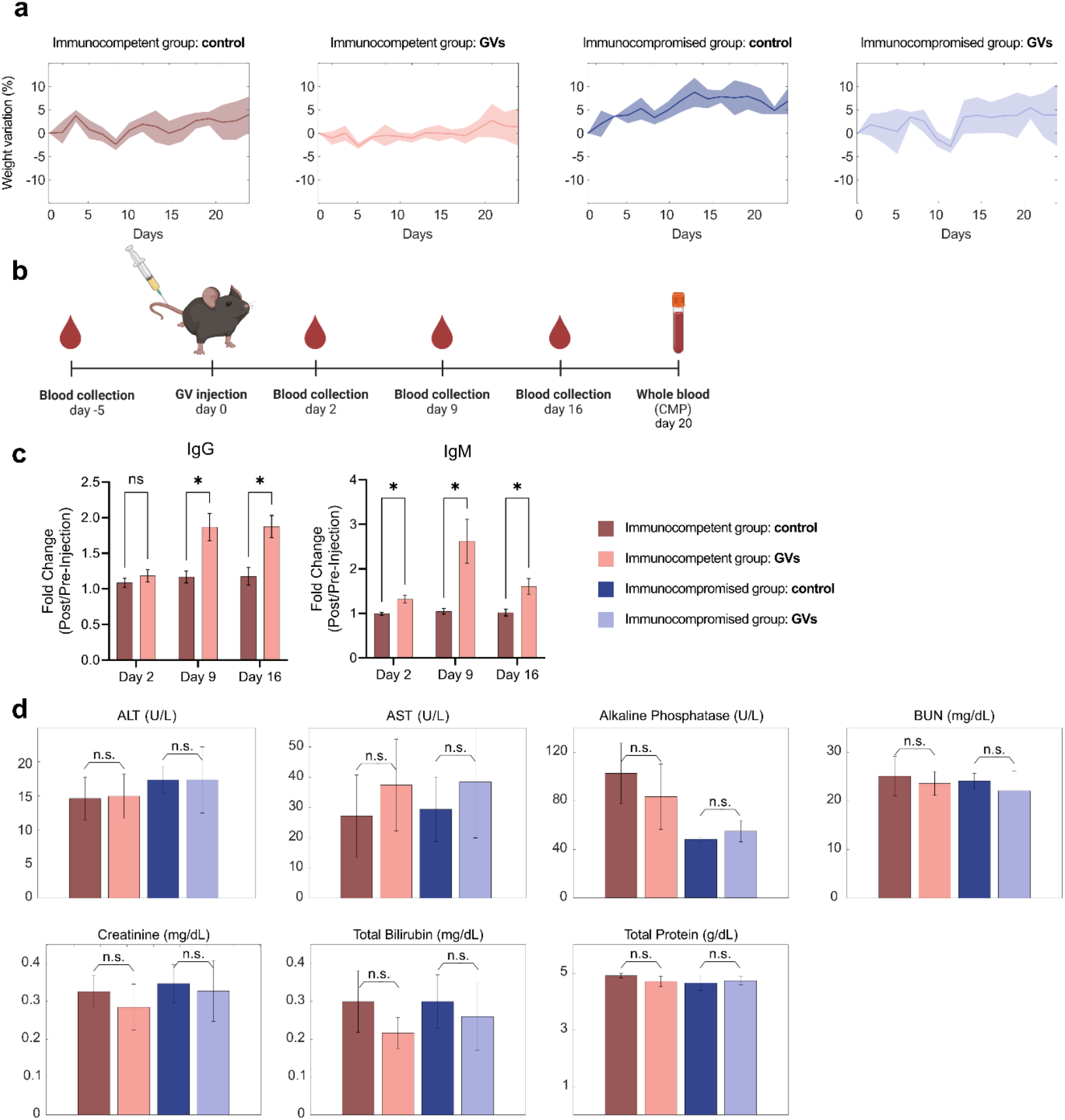
Body weight variation, serum immunoglobulin levels, and metabolic panel in mice infused with GVs. **a**, To characterize the clinical translatability of systemic GV injection, both immunocompetent C57BL/6 and immunocompromised NSG mice were IV infused with 150µl of 1.3 pmol purified GVs. Body weight was tracked over the subsequent days (sample sizes are N=4, N=6, N=5, N=5, respectively), demonstrating normal variability. **b**, Serum was collected 5 days prior to GV injection and on days 2, 9, and 16 post-injection. **c**, Bulk IgG and IgM levels at each of these time points were assessed in C57BL/6 mice. When compared with baseline, there was a significant increase in IgM levels after injection that peaked at around 9 days while IgG levels had a delayed increase, which corresponds with its slower production. Despite these increases, the mice displayed no hypersensitivity reaction and could be reinfused multiple times with no adverse effect. As noted, accelerated clearance and anti-PEG IgG and IgM is observed in other ultrasound contrast agents that are readministered, including FDA approved microbubbles^67^. **d**, A metabolic panel was run on day 20 after GV injection for both NSG and C57BL/6 mice. There was no significant increase in any of these biomarkers. Significance was determined with an unpaired t-test with Welch correction (alpha < 0.05). ALT = alanine aminotransferase, AST = aspartate aminotransferase, BUN = blood urea nitrogen.

**Figure S4:**
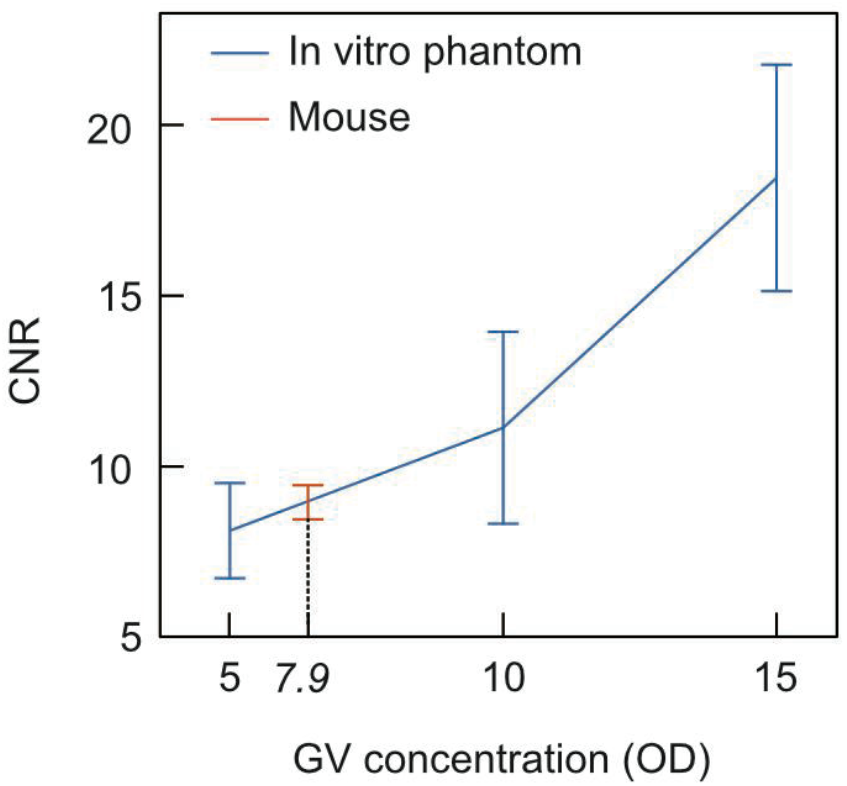
Comparison of contrast-to-noise ratio (CNR) across GV concentrations in vitro and in vivo. Phantom inclusions were prepared with varying GV concentrations to approximate those observed in GL261 tumors at day 7 post-implantation in immunocompetent mice. The graph indicates an average GV concentration of OD 7.9 in mice. Data for both phantom and mouse models were acquired using xAM imaging, with error bars representing standard deviations. Sample sizes were N=4 for each group

